# A modular transcript enrichment strategy for scalable, atlas-aligned, and clonotype-resolved single-cell transcriptomics

**DOI:** 10.64898/2026.02.06.703342

**Authors:** Mathini Vaikunthan, Anna C. M. Schoonen, Ioana-Teodora Lia, Jonathan Guo, José L. McFaline-Figueroa

**Affiliations:** Department of Biomedical Engineering, Columbia University, New York, NY, 10027, USA; Irving Institute for Cancer Dynamics, Columbia University, New York, NY, 10027, USA; Herbert Irving Comprehensive Cancer Center, New York, NY, 10032, USA

## Abstract

Existing targeted approaches either rely on per-gene barcoded probes and transcript tiling, which limit scalability, or forgo reverse transcription, precluding capture of highly variable transcripts. Here, we present targeted reverse transcription-linker (TRTL), a modular method that can be integrated into existing single-cell transcriptomic workflows to enable targeted readout of user-defined transcript panels, with minimal changes in protocol when scaling from tens to thousands of genes. Because TRTL retains reverse transcription, it also enables the capture of variable transcripts, such as TCR and BCR sequences. Applying TRTL and combinatorial indexing RNA-seq to the mouse brain, we show that carefully designed panels support robust alignment to existing reference atlases, enabling accurate cell type annotation and detection of cellular populations at low sequencing depths. Lastly, we combine TRTL with nuclear hashing-based multiplexing for a targeted-sci-Plex protocol and further demonstrate that targeted-sci-Plex can resolve dynamic T-cell fate trajectories following diverse activating exposures while concurrently profiling T-cell clonotypes.

## INTRODUCTION

Single-cell RNA sequencing (scRNA-seq) has revolutionized our ability to explore and catalog the transcriptomic landscape of diverse biological processes at cellular resolution, providing insight into development, organismal homeostasis, and the dysregulation that accompanies disease^1^. The last decade has resulted in significant gains in throughput, enabling the profiling of hundreds of thousands to millions of cells per experiment^2–6^. The coupling of these techniques to perturbation screens (e.g., Perturb-seq^7–11^, sci-Plex^12,13^) has further enriched atlas efforts. However, it remains a challenge to profile the large perturbation spaces that arise as cells respond to myriad endogenous and exogenous stimuli.

Designing single-cell experiments entails an explicit tradeoff between the number of cells, samples, or perturbations profiled and the depth to which each is characterized. As atlases expand to encompass time courses^14–17^, dose-responses^12,18,19^, variant-effects^20,21^, and combinatorial perturbations^22^, the limitations on fixed resources become restrictive. In these settings, a small number of mechanistically informative transcripts may be of primary interest, but are often sparsely represented in the resulting libraries. Existing strategies to enrich transcripts of interest attempt to address this imbalance. Biotinylated pull-down techniques post-library preparation can increase sensitivity, with high on-target read distributions, but are difficult to scale cost-effectively, as multiple probes per gene are needed to tile the entire transcripts (e.g., Parse Biosciences Gene Select kit^23^). Ligation-based probe capture of mRNAs, such as that implemented in the 10X Genomics Gene Expression Flex kit^24,25^, avoids reverse transcription and targets ∼50-bp stretches of a target transcript, yielding a modest yet scalable improvement in sensitivity^26^. However, these approaches do not query variable regions in transcripts, is susceptible to inaccurate quantification due to mispriming events when probes hybridize to off-target genes with high sequence similarity^27^, and are not as readily scalable as combinatorial indexing methods ^2,28,29^.

Recently, we and others developed transcript enrichment strategies for multiplexed combinatorial indexing-based single-cell RNA-seq, which, although increasing sensitivity, are not easily scalable over more than a handful of genes, as these strategies require a gene-specific RT barcoded plate^13,30,31^. Here, we address target scalability by creating a modular probe-generation strategy, the targeted-RT linker (TRTL), which we demonstrate can be coupled with single-cell combinatorial indexing RNA-seq. We further show that the TRTL module can be coupled to multiplex perturbation screens that rely on nuclear hashing^12,13^, resulting in a targeted version of our sci-Plex approach (targeted-sci-Plex). In addition, the TRTL module is designed to be integratable with existing combinatorial indexing single-cell workflows, such as Split-seq^5^ and SCI-LITE^30^, as well as single-cell rare-cell population targeting methods, such as PERFF-seq for targeted capture of cells and genes^32^ **(Supplementary Figure 1)**.

**Figure 1:**
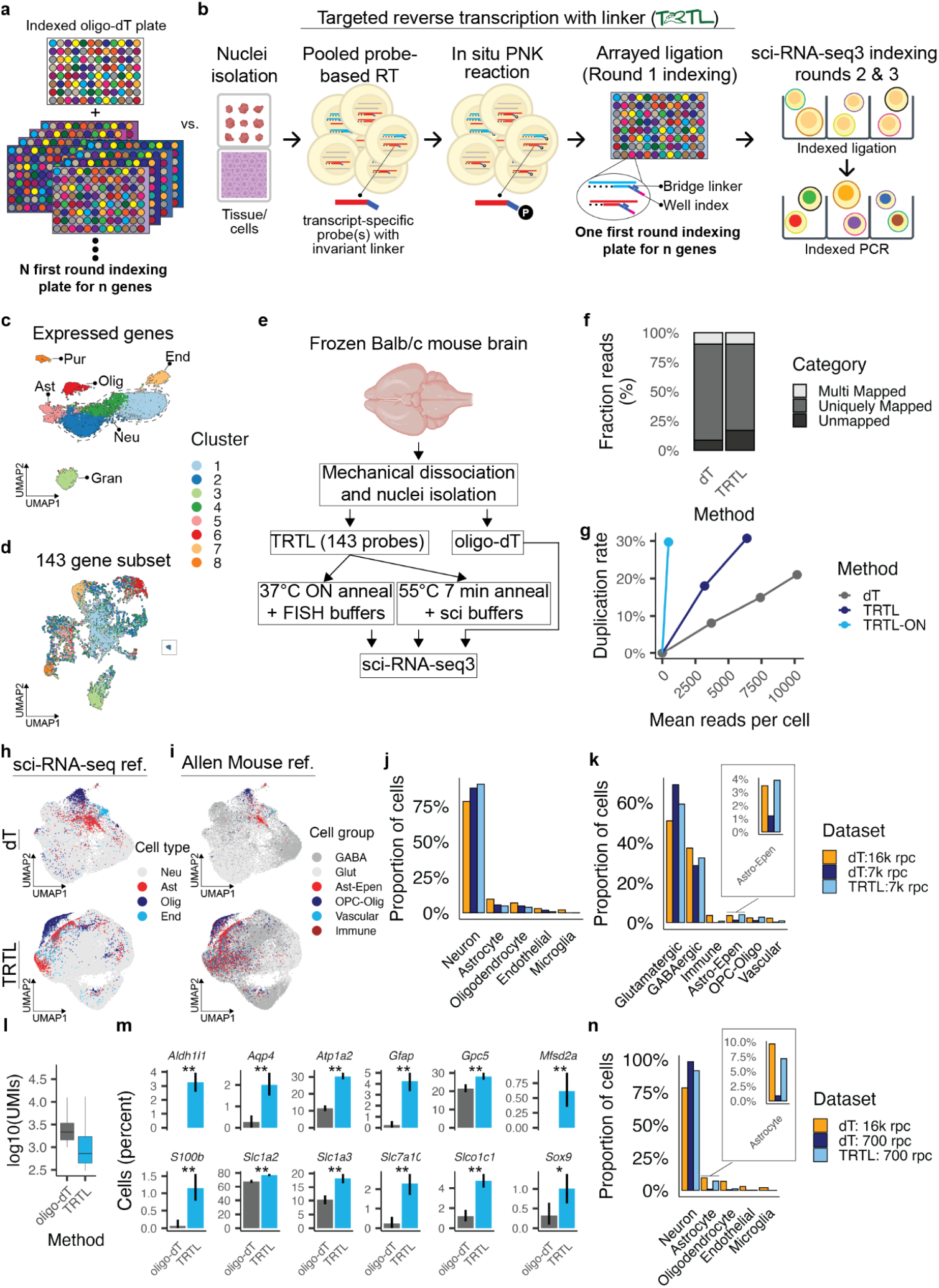
Targeted Reverse-Transcription with Linker (TRTL) enables cell type classification at low sequencing depths. **(a)** Existing targeted single-cell combinatorial indexing strategies use a set of targeted, barcoded oligonucleotides for each transcript profiled, which limits scalability across large numbers of genes. **(b)** Targeted reverse-transcription with linker approach highlighting pooled in situ reverse transcription across a panel of targeted genes, phosphorylation of target probe, and bridge ligation to append UMIs and the first round combinatorial index for compatibility with sci-RNA-seq3. **(c)** Batch aligned UMAP embedding of transcriptome-wide sci-RNA-seq3 profiling of the mouse brain, from which a panel of 143-gene targeting probes for cell type classification was designed. **(d)** Batch aligned UMAP embedding across a 143-gene subset chosen as target genes for broad cell type classification using TRTL. **(e)** Experimental overview of mouse brain nuclei isolation and processing for transcriptome-wide and TRTL protocols, including a 55°C 5 min anneal, similar to the standard sci-RNA-seq protocols, and a 37°C overnight (ON) anneal in more stringent single-molecule fluorescent in situ hybridization (FISH) buffers. **(f)** Mapping rates to the murine GRCm39 genome. **(g)** Duplication rate vs. reads per cell showing lowest complexity for the TRTL-ON condition, followed by TRTL and standard oligo-dT-based (dT) profiling. **(h-i)** Batch aligned UMAP embedding of downsampled (7,000 reads per cell [rpc]) for transcriptome-wide dT (top) and TRTL targeted (bottom) datasets. Color denotes the cell type classification using Seurat’s PCA-based anchor label transfer from the full transcriptome-wide reference generated by our sci-RNA-seq3 protocol (**h**) or to the Allen Mouse Brain Atlas (i). **(j-k)** Cell-type proportions annotated for downsampled dT and TRTL datasets at 7k rpc mapped to the full transcriptome-wide reference generated by our sci-RNA-seq3 protocol (j) or to the Allen Mouse Brain Atlas (**k**). **(l)** UMI counts for astrocyte assigned cells in the downsampled 7k rpc datasets (**m**) Percent of cells expressing more than one transcript for astrocyte marker genes in a dataset with equal information content per cell (normalized for UMI counts). Error bars indicate the 95% bootstrap confidence interval. Statistical significance of the difference between groups was determined by a non-parametric bootstrap test *\* p* < *0*.*05, ** p* ≤ *0*.*01*. **(n)** Cell type proportions as in **j** for a dataset downsampled to 700 rpc.

TRTL integrates a flexible probe-generation approach to simplify readout and increase the sensitivity of transcript detection with minimal changes in cost when scaling from 10s-100s-1,000s of genes of interest. By profiling cell types in the mouse brain, we demonstrate that careful selection of targeted transcripts enables alignment to existing reference atlases, facilitating robust cell typing and the detection of cellular populations. Because TRTL retains reverse transcription, it can interrogate variable transcripts that are inaccessible to approaches that rely on probe-dependent ligation. TRTL is also compatible with nuclear hashing and highly multiplex screens, enabling a targeted-sci-Plex approach that we demonstrate can map dynamic cell-fate transitions by profiling T-cell receptor (TCR) sequences to link transcriptional responses with clonotype information.

## RESULTS

Our targeted reverse-transcription linker (TRTL) approach provides a flexible, cost-effective, and scalable framework for targeted single-cell gene-expression analysis within the context of combinatorial indexing single-cell RNA-seq (sci-RNA-seq). In contrast to prior approaches that target a single gene with custom barcoded sets of oligonucleotides **(Figure 1a)**^13,30,31^, we devised a linker strategy that enables ligation of sequences necessary for the first round of cellular barcoding in sci-RNA-seq after reverse transcription (RT) across a pool of RT probes against a panel of genes of interest. In TRTL, we first perform RT in situ using a pool of 25-base pair (bp) oligonucleotide probes targeting a panel of transcripts of interest. Oligonucleotide probes contain a 15 bp 5’ invariant sequence upstream of the target-specific probe^5^. Targeted RT is performed across all cells in the experiment in bulk after which the 5’ ends of oligonucleotide probes are phosphorylated *in situ* with T4 polynucleotide kinase (PNK), and cells are distributed to multi-well plates. A bridge ligation reaction using a 30 bp linker oligonucleotide, then appends to each first strand cDNA a unique molecular identifier (UMI), the first combinatorial index, and a partial Illumina read 1 sequence **(Figure 1b)**.

### Mouse brain cell type identification and reference mapping with low read coverage

Landmark atlases and consortia, such as the Allen Mouse and Human Brain Atlas, the Human Cell Atlas, and Tabula Muris, have provided rich, annotated reference datasets of cell types across tissues and developmental stages^33–36^. These atlases, combined with evolving computational tools for reference mapping and annotation^37–39^, enable designs where users can achieve cell type identification and quantification of cell type proportions with targeted gene panels and cost-effective shallow sequencing. To determine the suitability of TRTL to generate cell typing data via a targeted workflow that maps to existing large-scale atlases, we first defined a minimal probe set for cell type identification based on transcriptome-wide sci-RNA-seq3 profiling of the mouse brain (Fig. 1c). We chose to distinguish the major cell types captured in unbiased (neurons, astrocytes, endothelial cells, oligodendrocytes, and microglia)^40^. As the neuronal population of the brain is comprised of hundreds of neuronal subpopulations, we selected a subset of target genes defining neuronal subtypes of interest from the literature, including Granule and Purkinje neuronal cells^40^, in addition to broad neuronal markers **(Supplementary Figure 2)**. In addition to canonical markers, we include genes that are significantly associated with cellular subsets from our sci-RNA-seq profiling of the mouse brain, defined as genes with the most positive and negative loadings for the first 10 PCs, as previously described^40^ **(Supplementary Figure 3a)**, resulting in a panel of 143 genes. To confirm that this 143-gene panel retained sufficient biological signal, we performed dimensionality reduction using only these genes and found substantial overlap in cell barcode membership between clusters annotated from the full transcriptome and those annotated from the 143-gene subset **(Figure 1d, Supplementary Figure 3b-c)**. The gene targeting sequences for our 143 gene panel were selected from the publicly available 10x Flex transcriptome probe set (10X Genomics), with 2-5 probes chosen for each gene, and a 24-25 base pair subset of each probe chosen based on GC content (40-60%GC) and a 3’ GC clamp to serve as our TRTL gene targeting probes **(Supplementary Table 1)**.

**Figure 2:**
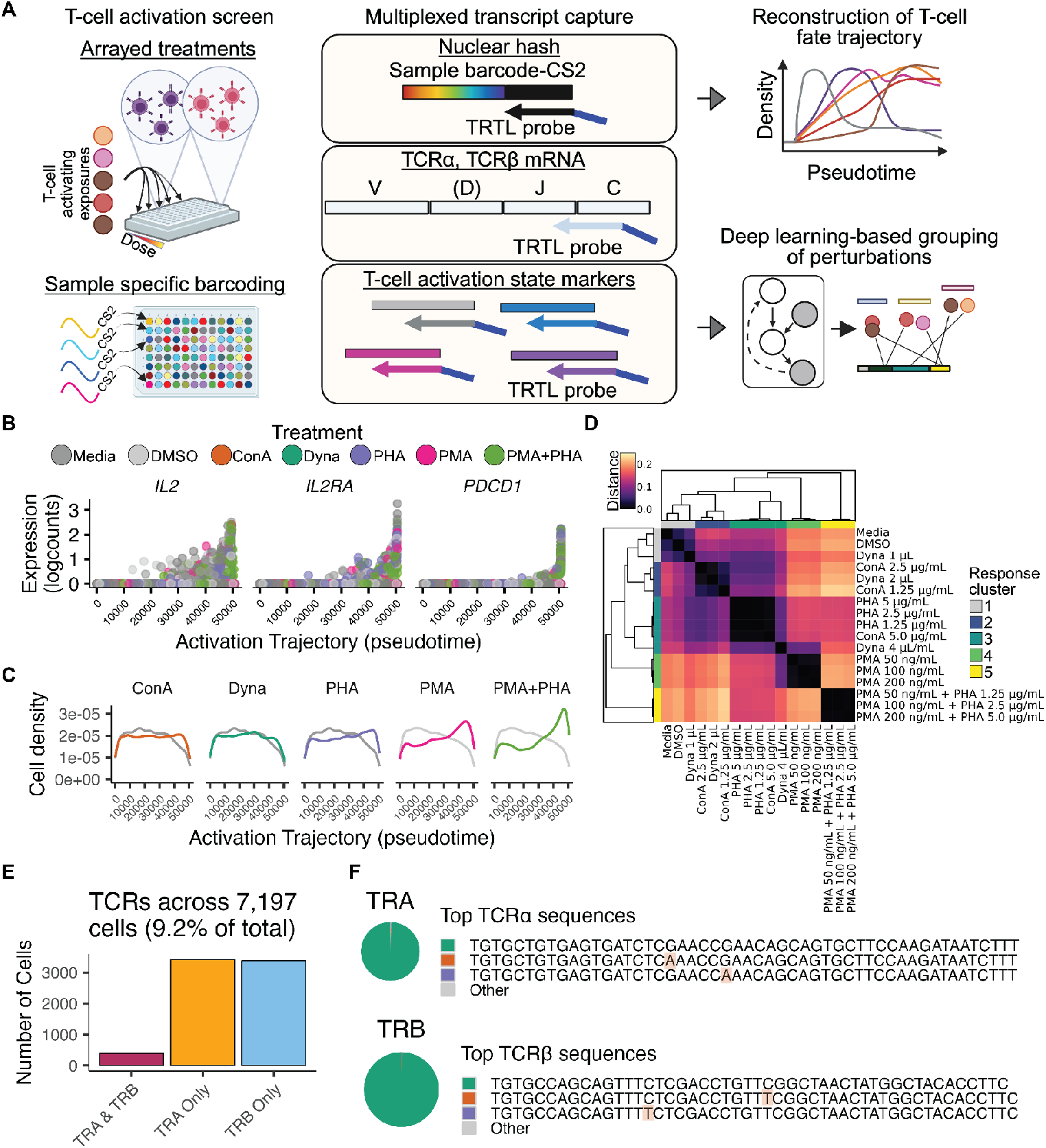
Targeted-sci-Plex maps dynamic cell fate trajectories and TCR clonotypes. **(a)** Targeted-sci-Plex profiling the effect of T-cell activation exposures on Jurkat T-cell fate. Jurkat cells across 51 distinct treatment conditions and replicates were labeled with sci-Plex hashes containing capture sequence 2 (10X Genomics), followed by TRTL-mediated enrichment of T-cell identity genes, T-cell activation markers, and TCRα and TCRβ constant regions. Targeted gene expression profiles were then used to infer a continuous response trajectory and a deep-learning based representation was used to cluster perturbations into modules with shared transcriptional effects. **(b)** Expression of T-cell activation genes across pseudotime defined by the top diffusion component (DC1) generated from the targeted sci-Plex experiment from Jurkat cells exposed to CD3/CD28 Dynabeads (dyna), concanavalin A (conA), phorbol myristate acetate (PMA), phytohemagglutinin (PHA), a combination of PMA + PHA, or DMSO and media controls. **(c)** Density plots of the distribution of cells exposed to the specified treatments along pseudotime for the top diffusion component across Jurkat cells exposed to dyna, conA, PHA, PMA, and PMA+PHA or DMSO and media controls. **(d)** Heatmap of the mean transcriptional distances between treatment conditions, estimated with MrVI. Hierarchical clustering of the pairwise distances reveal five distinct treatment clades based on transcriptional similarity and signaling pathway engagement (TCR vs. PKC activation). **(e)** Frequency and distribution of recovered TCRα and TCRβ transcripts across targeted-sci-Plex libraries. **(f)** Pie chart of TRTL recovered TCR sequences, showing high enrichment for the canonical Jurkat E6-specific TCRα and TCRβ variable regions (top sequence at right). Shaded areas in TCR sequences denote the bases at which the minor clones deviate from the top sequence.

Fresh-frozen mouse brains from Balb/C mice were isolated as previously described^18,41,42^ by pulverizing on dry ice, resuspending in nuclear EZ-lysis buffer, nuclei isolated by centrifugation, and fixed in formaldehyde. We then subjected nuclei to one of three gene quantification protocols: our standard transcriptome-wide sci-RNA-seq profiling approach, our TRTL targeted approach using standard sci-RNA-seq buffer conditions, and our TRTL targeted approach under an overnight probe annealing step in buffers derived from a single-molecule fluorescent in situ hybridization protocols (TRTL-ON)^43,44^ **(Figure 1e, Supplementary Figure 4a-c)**. The latter condition aims to examine strategies that decrease non-specific binding and its effects on atlas alignment.

The mapping of reads from our targeted libraries to the mouse Ensembl GRCm39 genome was high and comparable to our transcriptome-wide protocol (81.7% uniquely mapped reads in sci-RNA-seq, 73.3% uniquely mapped reads in TRTL) **(Figure 1f)**. As expected, sequencing saturation was reached rapidly for targeted libraries with the lowest complexity observed for the high-stringency smFISH-derived annealing condition **(Figure 1g)**. To compare performance across protocols, we downsampled libraries to a matched number of reads per cell (rpc). For our targeted protocol with reduced background annealing (TRTL-ON), we find a higher per-gene dropout rate relative to the transcriptome-wide and targeted preparations in sci-RNA-seq buffer (83/143 genes detected in TRTL-ON vs 120/143 genes detected in TRTL) **(Supplementary Figure 4h)**. This is consistent with FISH probe designs that often include dozens of probes to mitigate dropout while maintaining low background^45^, but may be too stringent given 2-5 probes per gene **(Supplementary Figure 4i-k)**. Therefore, we recommend moving forward using TRTL with sci-RNA-seq annealing conditions. We downsampled the dT and TRTL datasets to 7k rpc for comparison, and then used our higher sequenced (16k rpc) dT dataset (**Figure 1h**) or the Allen Mouse Brain Atlas (**Fig. 1i**) for label transfer. Aligning to our dT reference dataset using Seurat’s mapping and annotation workflow^46^, we identified well-resolved cellular neighborhoods for neurons, oligodendrocytes, astrocytes, and endothelial cells **(Supplementary Figure 4d-g)**. As expected, overall cell type proportions were highly concordant across the datasets at this sequencing depth (Pearson’s rho > 0.99 for dT and TRTL). However, we observed a dropout of cells that align with microglia when mapping to our sci-RNA-seq reference (**Fig. 1J**). Alignment to the richer Allen Mouse Brain Atlas (**Fig. 1k** and **Supplementary Figure 5d**) shows astrocyte and oligodendrocyte proportions are better maintained in our TRTL data compared to dT. We note a limitation in recovering microglia populations using both references (**Supplementary Figure 5e**), which we hypothesize can be remedied by additional microglia probes.

Consistent with targeted enrichment of cell type markers, the TRTL astrocyte and oligodendrocyte population had significantly less UMI diversity (**Fig. 1l, Supplementary Figure 5b**) yet higher expression of astrocyte- and oligodendrocyte-specific markers **(Figure 1m, Supplementary Figure 5e)**. Downsampling to just 700 rpc still allowed for the recovery of the neuronal and astrocyte populations in TRTL, whereas dT is unable to resolve the astrocyte population (**Figure 1n, Supplementary Figure 5f-h**). Defining probe sets that maximize enrichment, as observed for astrocytes, could, in turn, rescue additional cell populations at these very low sequencing depths.

### T-cell activation trajectories and TCR clonotypes with targeted-sci-Plex

Next, we sought to determine whether our targeted approach could be coupled to highly multiplex sample barcoding for single-cell RNA-seq^12^ to enable large-scale perturbation screens with targeted readouts. As a proof-of-concept, we profiled T-cell activation in the Jurkat cell line using multiple treatments at varying doses, providing an opportunity to test whether targeted transcriptome assays can recover biologically meaningful trajectories while capturing variable T-cell receptor (TCR) transcripts that are not accessible in targeted approaches lacking a reverse transcription step. Jurkat cells were exposed for 24 hours to (1) CD3/CD28 functionalized beads (dynabeads), concanavalin A (conA), phorbol myristate acetate (PMA), phytohemagglutinin (PHA), and a combination of PMA+PHA at three doses centered on standard activation conditions (dynabeads^47^: 1 μL, 2 μL, 4 μL; conA^48^: 1.25 μg/mL, 2.5 μg/mL, 5 μg/mL; PHA^49^: 1.25 μg/mL, 2.5 μg/mL, 5 μg/mL; PMA^50^: 50 ng/mL, 100 ng/mL, 200 ng/mL; PHA and PMA Combination^51^: 1.25 μg/mL and 50 ng/mL, 2.5 μg/mL and 100 ng/mL, 5 μg/mL and 200 ng/mL) alongside media and DMSO controls. The 24-hour timepoint was chosen based on previous studies of increased protein expression of activation-associated factors (*IL-2*^*52*^ and *PD-L1*^*51*^). After treatment, we isolated nuclei and uniquely barcoded nuclei from each exposure with sci-Plex hashes containing capture sequence 2 (CS2)^10^ instead of the polyA capture sequences used in our current^12,18^ transcriptome-wide multiplex profiling protocol. We then applied TRTL and sci-RNA-seq3 using a panel of 8 genes targeting general markers of Jurkat/T-cells (*PTPRC, CD8, CD4, CD3G*), T-cell activation markers (*IL2, IL2RA, PDCD1, CD69*), and probes targeting the constant region of TCRα and TCRβ ^53^, designed to capture a ∼250-bp fragment around the variable region (**Figure 2a, Supplementary Figure 7a**). Following filtering based on UMI and hash UMIs, we recovered a median of 7077 cells per treatment, and 2315 cells per treatment-dose condition.

Dimensionality reduction with diffusion maps^54,55^ recovered a trajectory consistent with T-cell activation with treatment- and dose-dependent **(Supplementary Figure 6a-c)** increases in *IL2, IL2RA*, and *PDCD1* along the dominant diffusion component (**Figure 2b**). Comparing the relative proportion of cells from each condition along the trajectory revealed the extent to which each exposure shifted cells away from the baseline state defined by the media and DMSO controls (**Figure 2c, Supplementary Figure 7b**). Dynabeads and conA treatment produced only modest increases in the fraction of T-cells occupying terminally activated states, whereas PHA-treated cells appeared to enrich at the end of the activation trajectory. PMA alone induced strong activation and accumulation at high pseudotime values, which were further amplified by PMA+PHA co-treatment. To quantitatively compare transcriptional responses across treatments, we used multi-resolution variational inference^56^ (MrVI) to compute sample-sample distances across each condition and group conditions by transcriptional similarity (**Figure 2d**). The MrVI analysis using probe genes and 500 HVG genes identified five major clades: Cluster 1, composed of DMSO and media controls and low dose dynabead stimulation; Cluster 2, composed of intermediate dynabeads and conA exposure; Cluster 3, containing PHA-treated cells, with high dose dynabeads and conA clustering with PHA; Cluster 4, composed of PMA-treated cells; and Cluster 5, composed of PHA + PMA co-treated T-cells. This hierarchy is consistent with differences in pathway engagement across our exposures. Dynabeads, conA, and PHA initiate activation via the TCR/CD3 complex and upstream signaling, whereas PMA directly activates the downstream effector Protein Kinase C (PKC) **(Supplementary Figure 8b)**, with PMA+PHA providing co-stimulatory input that interestingly is transcriptionally distinct from the component inputs, as also shown in the MrVI z-latent space capturing sample effects **(Supplementary Figure 7c)**. The distinct activation profile of PMA+PHA is recovered using only probe genes as input for MrVI **(Supplementary Figure 8a)**. This suggests that highly stringent targeted data—even without background genes for normalization—can effectively group large-scale combinatorial screens.

Within this multiplexed targeted-sci-Plex experiment, TRTL-mediated capture of the TCR constant region recovered the expected Jurkat E6 TCRα and TCRβ sequences in ∼9.2% of cells without dedicated TCR enrichment (**Figure 2e**). The recovered TCR sequences were highly enriched for the expected Jurkat E6 TCRα and TCRβ sequences (**Figure 2f**). Overall, our results demonstrate that targeted-sci-Plex can jointly resolve perturbation responses and TCR clonotypes within a single, targeted combinatorial indexing workflow.

TRTL enables low-sequencing-depth cell type transfer and highly multiplexed TCR and dynamic state shift capture. Our approach enables flexible atlasing and large-scale screens where key marker genes are highly informative for cell state transitions while reducing sequencing requirements and increasing sensitivity. In principle, TRTL is also compatible with recent technologies for targeting rare cell types (PERFF-seq^32^) in multi-tissue/organism experiments, enabling cell- and gene-targeting in highly cell-type- and cell-state-specific screening.

## Data and code availability

Raw and processed data will be available at NCBI GEO upon publication. The code necessary to reproduce the analyses in this study can be found at github https://github.com/mcfaline-figueroa-lab/targeted-sci-Plex. The code to run MrVI can be found at https://scvi-tools.readthedocs.io/en/latest/tutorials/notebooks/scrna/MrVI_tutorial.html.

## Supplementary Information

Supplementary Information file contains

Supp. Fig. 1. Comparison of gene-targeted methods and compatibility of TRTL, in principle, with single-cell methodologies

Supp. Fig. 2. Cell type and cell lineage specific marker expression in mouse brain samples subjected to transcriptome-wide combinatorial indexing single- cell RNA-seq profiling Supp. Fig. 3. Mouse brain panel selection

Supp. Fig. 4. Transcriptome-wide oligo-dT vs TRTL and TRTL-ON sci-RNA-seq profiling Supp. Fig. 5. TRTL 7k and 700 reads per cell (rpc)

Supp. Fig. 6. Recreating a continuous T-cell activation trajectory with diffusion maps Supp. Fig. 7. Deep learning based classification of T-cell activating exposures from a targeted-sci-Plex readout

Supp. Fig. 8. Classification of activating agents based on only probe genes

Supp. Table 2. Overall sequencing and subsampling summary for mouse brain

Supp. Table 1 can be found as a separate spreadsheet file and contains final mouse brain panel, number of probes per gene in the mouse brain studies, final Jurkat panel, number of probes per gene in the Jurkat studies, oligos used in studies

## Acknowledgements

We thank the members of the McFaline-Figueroa, Simunovic, Correa, Azizi laboratories, and P.T. Ravindran for their helpful discussions during the development of this study. We thank the Correa laboratory, particularly A. Margaronis for providing the Jurkat, E6-1 clone. We thank E. Azizi, N. Hou and P.T. Ravindran for their helpful comments on the manuscript. This work is supported by grants to J.L.M.-F from the NIH (R35HG011941), the NSF (2146007), and by an Allen Distinguished Investigator Award, a Paul G. Allen Frontiers Group advised grant of Allen Family Philanthropies. The authors would like to thank Erin Bush and the JP Sulzberger Columbia Genome Center for their support with next-generation sequencing; this research was funded in part through the NIH/NCI Cancer Center Support Grant P30CA013696 and used the Genomics and High Throughput Screening Shared Resource. **Author contributions**: J.L.M.-F conceived the project and provided overall supervision of the study; M.V. designed experiments; M.V. and J.L.M.-F. analyzed and interpreted the data; M.V., A.S., I.L., and J.G. performed experiments or aided in the analysis. M.V. and J.L.M-F wrote the manuscript with input from all authors.

## METHODS

### Experimental approaches

#### Tissue

Mouse brain tissue samples were purchased from The Jackson Laboratory. Specifically, brains isolated from 6-8 week BALB/cJ 000651 female mice, flash frozen with stem (no fixative, perfused) and stored at -80C until processing.

#### Nuclei isolation and fixation from brain tissues

Nuclei isolation from mouse brain samples, hashing, and fixation were adapted from Martin et al.^41^ and Sziraki et al.^42^ Briefly, tissues were ground to a powder on dry ice before lysing in a 50 mL conical tube with 4mL EZ Lysis Buffer (Sigma) supplemented with 1% diethyl pyrocarbonate (Sigma), 0.1% SuperaseIn RNase Inhibitor (Thermo). After lysis, nuclei were strained through a 40 μm cell strainer into EZ lysis buffer and nuclei pelleted. The nuclei were fixed by addition of formaldehyde in Dulbecco’s phosphate-buffered saline (DPBS) to a final concentration of 1% formaldehyde and incubated on ice for 10 minutes. Nuclei were pooled into a reagent reservoir and transferred to a 50mL conical tube for centrifugation at 650 x *g* for 5 minutes at 4°C. Supernatant was removed, and nuclei washed with nuclei suspension buffer (NSB; 10 mM Tris-HCl, pH 7.4, 10 mM NaCl, 3 mM MgCl2, 1% Superase RNA Inhibitor (Thermo Fisher), 1% 20mg/mL Recombinant Albumin (New England Biosciences)). Finally, nuclei were resuspended in NSB, slow-frozen in 10% DMSO, and stored at −80°C until sci-RNA-seq processing.

#### Cell lines and cell culture

Jurkat, Clone E6-1 were purchased from ATCC. Cells were cultured in RPMI media (ThermoScientific) supplemented with 2mM L-glutamine (FisherScientific), 10% fetal bovine serum, and 1% penicillin/streptomycin (P/S, ThermoScientific) as specified by ATCC. Cells were cultured in a 37 °C tissue culture incubator with 5% CO2. The cell line was used within 10 passages and routinely tested for mycoplasma contamination.

#### Jurkat activation screen perturbation procedure and materials

Jurkat cells were plated at a density of 8 x 10^4^ cells per well in 200 µL of media in 96-well v-bottom plates and allowed to acclimate overnight. Gibco Dynabeads Human T-Activator CD3/CD28 (Cat. No. 111.31D) was purchased from ThermoFisher Scientific; 150 µL were washed in DPBS with 0.1% recombinant albumin (DPBS w/ 0.1% BSA) and resuspended in 150 µL media. A volume of 1 µL, 2 µL (manufacturer recommended), or 4 µL was added to a final volume of 200 µL. eBioscience Phytohemagglutinin-L (PHA-L) Solution (00-4977-93) (500x, 1.25mg/mL) and Concanavalin A (Con A) Solution (Cat. No. 00-4978-93) (500X, 1.25mg/mL) were purchased from ThermoFisher Scientific and diluted 5-fold in media to make a 100x solution. A volume of 1 µL, 2 µL (manufacturer recommended), or 4 µL of 100x solution was added to a final volume of 200 µL. Phorbol 12-Myristate 13-Acetate (PMA) (5005820001) 10 mM in DMSO was diluted to 1 mg/mL in DMSO. This was diluted 100-fold in media to yield the 100x solution. A volume of 1 µL, 2 µL (manufacturer recommended), or 4 µL of 100x solution was added to a final volume of 200 µL. DMSO concentration was normalized to 0.2%v/v DMSO for all the PMA wells and the DMSO control wells. Three well replicates were present for each condition, and nine well replicates were present for each control. Nuclei were harvested 24 hours post-exposure.

#### Nuclei hashing with hash of Jurkat screen

Nuclei hashing and fixation procedures were adapted from McFaline-Figueroa et al.^13^, Srivatsan et al.^12^ and Sziraki et al.^42^ Briefly, suspension Jurkat cells were dissociated by pipetting. Upon washing with ice-cold 1X PBS, cells were lysed with EZ Lysis Buffer (Sigma) supplemented with 1% diethyl pyrocarbonate (Sigma), 0.1% SuperaseIn RNase Inhibitor (Thermo Scientific), and 5000 fmol of hashing oligo. After lysis, nuclei were fixed by adding 1.25% formaldehyde in 1X PBS (final well concentration 1% FA) and incubated on ice for 10 minutes. Nuclei were pooled into a reagent reservoir and transferred to a 50mL conical tube for centrifugation at 650 x *g* for 5 minutes at 4°C. Supernatant was removed from the nuclei pellet, and nuclei were washed once with nuclei suspension buffer (NSB; 10 mM Tris-HCl, pH 7.4, 10 mM NaCl, 3 mM MgCl2, 1% Superase RNA Inhibitor [Thermo Fisher], 1% 0.2mg/mL Ultrapure BSA [New England Biosciences]). Nuclei were resuspended in NSB, slow-frozen in 10% DMSO, and stored at −80°C until sci-RNA-seq processing.

#### Targeted Reverse Transcription with Linker Probe Design

We selected gene targeting sequences from the publicly available 10x Genomics Fixed RNA profiling (Flex) mouse and human transcriptome probe sets. The selection process was designed to repurpose the optimized sequences into 25-bp single-stranded oligonucleotides for targeted RT. We identified probe pairs from the 10x Genomics Flex v1.0 reference corresponding to the chosen gene set for each experiment (either Chromium Human Transcriptome Probe Set v1.0.1 GRCh38-2020-A for human or Chromium Mouse Transcriptome Probe Set v1.0.1. Each 10x Flex probe consists of two 25bp segments (a 5’ probe and 3’ probe) that together make a 50bp segment for hybridization to a transcript. We chose either the 5’ or 3’ probe based on a GC content between 40-60%, and a GC clamp (terminal G or C at 3’ end), or presence of a G or C within the last 5 nucleotides at the 3’ end. We then append a constant invariant 5’ handle of the form: 5’-AGGCCAGAGCATTCG-3’

which serves as the site for the subsequent bridge ligation of the first barcode, UMI, and partial R1.

The final gene probe takes the form:

5’-AGGCCAGAGCATTCG-[25 bp or longer gene targeting probe]-3’

The final hash targeting probe for CS2 capture takes the form:

5’-AGGCCAGAGCATTCGCCTTAGCCGCTAATAGGTGAGC-3’

Gene-targeting oligos were ordered either as a pool or as individual probes (IDT). For pools, oligos were ordered without 5’ phosphorylation as this substantially increases the per-probe cost. Instead, we perform *in situ* phosphorylation of the probes after RT using PNK kinase (New England Biolabs). For the mouse brain panel, 143 genes with ∼3 probes per gene (428 total) were ordered as two oPools from IDT, with 50 pmol per oligo. For the Jurkat activation screen, 8 genes and 2 targets for TCR constant regions were chosen with phosphorylation and ordered as individual single-stranded DNA (25nmol) with 5’ modification, as this was a small set of probes.

### Targeted Reverse Transcription with Linker Sci of mouse brain nuclei

#### Nuclei preparation

Formaldehyde-fixed mouse brain nuclei (500,000–2,000,000) were thawed at 37 °C and pelleted at 700g for 5 min at 4 °C. Nuclei were resuspended in 100 µl NSB. For permeabilization, a mixture of 390 µl NSB and 10 µl 10% Triton X-100 was added and incubated on ice for 3 min. Nuclei were pelleted at 700g for 5 min, resuspended in 400 µl NSB, and sonicated using a Bioruptor (low power mode, 12s). The suspension was filtered through a Flowmi cell strainer, pelleted again at 500g, and resuspended in 100 µl NSB for quantification.

#### Parallel processing of TRTL gene-specific reverse transcription and standard sci-RNA-seq3

The nuclei were divided into three experimental arms performed in parallel: a standard sci-RNA-seq preparation, and the TRTL workup with either TRTL or TRTL-overnight (ON).

For the TRTL-ON, annealing occurred the night prior to processing with TRTL sci-RNA-seq3. Single-molecular fluorescence in situ hybridization (smFISH) hybridization and wash buffers were prepared as described previously^44^. Formaldehyde-fixed nuclei after sonication (500,000–2,000,000) were pelleted at 800 x *g* and resuspended in 80 µl Hybridization Buffer (10% w/v dextran sulfate, 10% v/v formamide, and 2X Saline-Sodium Citrate Buffer [SSC]). 20 µl of the 50 µM gene-specific primer pool was added, and the reaction was incubated overnight at 37 °C. To remove unhybridized oligos, 900 μl of Post-Hybridization Wash Buffer (10% v/v formamide, 2X SSC) was added directly to the sample. 175 μl was used for resuspension before transferring to a 1.5 ml microcentrifuge tube to reach a final volume of 900 μl. The suspension was incubated at 37 °C for 10 min in a thermomixer. Nuclei were then pelleted at 850 x g for 5 min at room temperature. The supernatant was carefully aspirated to avoid disturbing the pellet, and nuclei were resuspended in NBB for downstream processing.

For the TRTL gene-specific arm, pooled nuclei (1500-2500 nuclei per μl) in 180 μL NBB per well were combined with 32 µl of 10 mM dNTPs and 32 µl of 25 µM gene-specific primers (100 pmol total). Annealing was performed at 55 °C for 7 min for TRTL.

The washed nuclei for TRTL and TRTL-ON processed nuclei were resuspended in NBB at a concentration of 1,500-2,500 nuclei per µl. For each targeted sample (TRTL, TRTL-ON), a reverse transcription master mix was prepared by combining 80 µl 5X RT Buffer (Thermo Scientific), 20 µl nuclease free water, 20 µl Maxima H Minus reverse transcriptase (Thermo Scientific), and 20 µl SuperaseIn RNase Inhibitor (Thermo Scientific); 128 µl of this mix was added to the for a total volume of 372 µl per sample condition. The reaction was split up into 4 wells with 93 µl per well, and reverse transcription was carried out in a thermocycler using the following ramped program: 4 °C for 2 min, 10 °C for 2 min, 20 °C for 2 min, 30 °C for 2 min, 40°C for 2 min, 50 °C for 2 min, and 55 °C for 15 min^18^.

For the sci-RNA-seq-3 arm, RT was performed as described previously^18,20,57^. Briefly, sonicated nuclei were resuspended in 400 μl at a concentration of 1000-2000 nuclei per μl, and divided into 16 wells with 21 μl of nuclei in NBB distributed into each well with with 2µL 10mM dNTP, 2µL 100 µM indexed oligo-shortdT primers, 2µL 100 µM indexed random hexamer primers, and 14µL of a reverse transcription master mix consisting of 14.29% nuclease free water, 14.29% SuperaseIn RNase Inhibitor, 57.14% 5X RT Buffer, and 14.29% Maxima H Minus reverse transcriptase, which are the same ratios used in the TRTL procedure. Reverse transcription was carried out in a thermocycler with the same ramped program used for TRTL, with the corresponding volume per well set on the thermocycler.

#### In situ probe phosphorylation and bridge ligation for TRTL and TRTL-ON samples

Post-RT, TRTL and TRTL-ON nuclei were washed once in NBB and subjected to phosphorylation. A 660 µl reaction was prepared containing 66 µl T4 DNA ligase buffer (NEB), 33 µl T4 PNK enzyme (10,000 units/ml) (NEB), and 561 nuclei in NBB. The reaction was incubated at 37 °C for 30 min. Following PNK treatment, nuclei were pelleted and resuspended in 115 µl NBB. Nuclei were distributed (3 µl per well) and 1 µl of a 50 µM bridge oligo of the form:

5’ CGAATGCTCTGGCCTCTCAAGCACGTGGAT 3’

was added, followed by a 15-minute thermal ramp to 55 °C. Combinatorial barcodes were introduced by adding 2 µl of 25 µM Barcode 1 oligo of the form:

5’ CAGAGCNNNNNNNN-[10bpBarcode1]-ATCCACGTGCTTGAG 3’

with ligation performed at 25 °C for 1 hour.

#### Ligation of the second round indexing barcode

Following the initial barcoding, all experimental arms were treated identically for the next two barcoding rounds. Nuclei were pooled and resuspended in 2.03 ml NSB per 96-well plate. The suspension was distributed into a new 96-well plate at 19.7 µl per well. To each well, 8 µl of 100µM indexed ligation primer of the form:

5′-GCTCTG-[9bp-or-10bp-barcode-A]/ideoxyU/ACGACGCTCTTCCGATCT[reverse-complement -of barcode-A]-3’

was added. A ligation master mix was prepared by combining 288 µl T4 buffer and 192 µl T4 ligase (400,000 units/ml) (NEB), and 4.2 µl of this mix was distributed to each well. The ligation reaction was performed at 25 °C for 1 hour.

#### On-plate second-strand synthesis and tagmentation

Nuclei were pooled, washed twice in 1 ml NBB, and resuspended in 500 µl NBB. After quantification, nuclei were distributed into 96-well plates at 5 µl per well and stored at -80 °C. Upon thawing, 5 µl of second-strand synthesis mix (New England Biolabs) 330 µl elution buffer, 146.7 µl second strand buffer, and 73.3 µl enzyme) was added per well and incubated at 16 °C for 3 hours. Tagmentation was performed by the addition of 10 µl of tagmentation mix composed of 1.1 ml tagmentation buffer (20 mM Tris-HCl, 10mM MgCl_2_, 20% v/v dimethylformamide) and 2.75 µl of in-house N7 adaptor-loaded Tn5 (Macro Lab UC Berkeley) per well at 55 °C for 5 min. The reaction was quenched with 20 µl DNA-binding buffer (Zymo Research).

#### Library purification and PCR amplification

Purification was performed using 40 µl CleanNGS beads (Bulldog Biosciences) per well. Libraries were eluted in 17 µl buffer, and 16 µl was used for PCR with 2 µl P5 primer, 2 µl P7 primer, and 20 µl NEBnext master mix (New England Biolabs). PCR conditions were as follows: 72 °C for 5 min, 98 °C for 30 s, followed by 12 to 15 cycles of 98 °C for 10 s, 66 °C for 30 s, 72°C for 30 s, and 72 °C for 5 min.

#### Final purification and quantification

Pooled PCR products were purified with 0.7X CleanNGS beads (Bulldog Biosciences) and eluted in 100 µl of Buffer EB (Qiagen). For hashed samples, the supernatant was recovered, and a 1X cleanup was performed using 150 µl CleanNGS beads per 500 µl of supernatant. Library concentration and size distribution were quantified using the Qubit 4 fluorometer (Thermo Fisher) and TapeStation D1000 (Agilent), respectively.

### Targeted Reverse Transcription with Linker sci-Plex of Jurkat screen

The same method for TRTL was used for the hashed sci-Plex sample of the Jurkat screen, except in-situ PNK did not take place as the probes were ordered with modification: 5’ Phosphorylation.

### Data preprocessing and generation of count matrices

#### Sequencing, fastq generation, and barcode whitelist matching

For the mouse brain profiling experiments, sequencing was performed either on Element Bioscience AVITI (2x75 paired end) or using Novogene’s standard workflow on the Illumina NovaSeqX Plus platform standard workflow (2x150 paired end) and final reads were trimmed to R1:35, R2: 101, I1:10, I2:10. For the Jurkat profiling experiment, Element Bioscience AVITI 2x150 was used to reach R2: 251 (R1:35, I1:10, I2:10 stayed the same).

Raw reads were transformed to fastq using bases2fastq v2.20.0.422 or bcl2fastq 1.5.0.962525890 for Element AVITI and Illumina NovaSeqX, respectively. A custom data processing pipeline adapted from Srivatsan et. al was used to process fastq data into a single-cell count matrix. First, reverse transcription and ligation barcodes were assigned to reads with a mismatch allowance of 1bp. Reads were assigned to either random hexamer or oligo-shortdT or oligo-linker based on the whitelist from these primer sequences. Random hexamer and oligo-shortdT reads once correctly mapped to their barcodes were collapsed into a single folder for processing each PCR well. After index assignment, polyA sequences were trimmed using TrimGalore version 0.6.10 and CutAdapt version 2.6.

#### STAR alignment

Upon polyA trimming, reads were aligned to mouse GRCm39, or human GRCh38 respective of the experiment using STAR version 2.7.9a. The Log.out.final summary mapping statistics output was collated to map the rate of Multi Mapped, Uniquely Mapped, and Unmapped reads between the TRTL and dT experiments.

#### Generation of count matrix and quality control

Finally, aligned reads were filtered for quality and duplicates and assigned to genes using bedtools version 2.26.0 as described previously. The resulting unique read assignments were aggregated by UMI and collapsed by cell and gene, yielding a raw sparse count matrix. The raw sparse count matrix, cell annotations, and gene annotations were used to make a cell_data_set (cds) object using the R package *monocle3*^*2,58,59*^. Cell barcodes for all experiments (including TRTL, TRTL-ON and targeted-sci Plex) were filtered using a UMI cutoff determined visually with a kneeplot of cell rank by UMI count, which retained its characteristic shape **(Supplementary Figure 4)**.

For targeted-sci-Plex, hash assignments were determined similarly to described previously (Srivatsan, McFaline-Figueroa). Hash barcodes were assigned if the read was adjacent to the CS2 sequence, and if the barcodes matched a whitelist with a mismatch allowance of 1bp. Duplicate hash reads were rolled up by UMI and collapsed into hash assignment counts by cell. Hashes were assigned to cells by two criteria: (1) having ≥ 3 hash UMIs and (2) a ratio of the cell’s top hash UMI to second best hash UMI of 1.5. This resulted in 50774 cells with hash out of 78129 cells for the targeted-sci-Plex.

The *monocle3* package was used to manipulate, batch align, and visualize the data throughout.

#### Downsampling reads

Downsampling was performed in the data analysis pipeline by processing the STAR-aligned bam files (including unmapped reads) via samtools view -b -s, which downsamples reads according to a target fraction of the original reads. Fractions were approximately calculated to achieve a final reads per cell of ∼7000 reads per cell, or ∼3500 reads per cell when inspecting the complexity (duplication rate at various reads per cell) of targeted and transcriptome-wide libraries. Fractions were approximately calculated to achieve 700 rpc for the very low sequencing depth comparison (**Supplementary Figure 6**). Once downsampled, the steps of the generation of count matrix and quality control were repeated for each sample.

### Probe set design

To capture the cellular diversity of the murine brain, a targeted panel of 143 genes was curated. This selection was informed by several criteria, including canonical markers for general neuronal and non-neuronal identity, as well as for distinct neuronal subpopulations such as Purkinje and Granule neurons. The panel also incorporated cluster-specific markers and genes exhibiting the highest positive and negative loadings across the first 10 principal components derived from preliminary whole-transcriptome analysis determined using Seurat. To ensure that the reduced feature set retained sufficient biological signal, dimensionality reduction was applied to the 143-gene subset. Clustering and UMAP projections were executed within the aligned integrated space of the mouse brain samples to account for biological and technical batch effects. We then performed clustering on this 143-gene space and compared the proportions of cells in the original clusters with those in the new clusters, showing that the targeted gene space preserved both the global and local structures of the original whole-transcriptome clusters **(Supplementary Figure 3)**.

For mouse brain and T-cell activation profiling, targeting sequences were derived from the 10x Genomics Flex transcriptome probe set to utilize pre-validated, high-specificity sequences. For the TRTL gene targeting probes, a 24-25 base pair subset was selected from these sequences, optimized for a GC content of 40-60% and specific 3’ terminal stability parameters to ensure robust hybridization kinetics.

### Aligning of targeted and downsampled single-cell profiles to reference atlases

#### Seurat Label Transfer

Label transfer was performed using Seurat. First, counts in each experimental replicate were normalized using SCTransform and aligned using the JointPCAIntegration function, which creates an integrated dimensional reduction space based on the joint PCA of the two experiments. The first 15 dimensions of the joint PCA reduction from the full dT dataset were used to find transfer anchors to the query space (downsampled dT and TRTL datasets), using the chosen probe genes and variable feature genes from the SCTransform normalized counts. The anchors were used to transfer the cell type labels determined on the full dT dataset to the downsampled dataset.Pearson correlation coefficients were calculated by comparing the proportions of each identified cell type between the 7k and 16k rpc datasets for the standard and TRTL conditions respectively.

#### Allen Brain Atlas

Cell type annotation was performed using MapMyCells from the Allen Brain Atlas, a tool that maps single-cell transcriptomic data to reference taxonomies. Input data consisted of a cell-by-gene matrix in csv format with gene symbols or identifiers. Mapping was performed via the MapMyCells web interface using the 10x Mouse Brain reference taxonomy and the hierarchical mapping algorithm. Mapping results were integrated with cell metadata for downstream visualization and comparison using R scripts in GitHub. Cell type annotations were transferred as “class_name”. Cell type groupings were created for GABA and Glut from the class_name by extracting the “GABA” or “Glut” from the “class_name”.

### Percent cell positive plots using normalized UMI

To visualize marker gene expression across the dataset, UMI counts were downsampled to a target median depth to equalize the contribution of individual cells and minimize library-size-related biases. This process was performed by calculating a cell-specific downsampling factor, defined as the ratio of the target median UMI count to the total observed UMI counts for each cell, with a maximum factor capped at one. For cells exceeding the target depth, UMIs were stochastically downsampled without replacement to match the calculated fraction, following the logic implemented in the Seurat *SampleUMI* framework. Cells were then plotted using *percent_cells_positive* from the *monocle3* package.

### Bootstrap-Based differential expression testing

To assess the statistical significance of the difference in expression frequency between groups, a non-parametric Bootstrap Hypothesis Test was performed using the *boot* R package. For each gene, the dataset was resampled with replacement over 100 iterations (bootstrap_samples = 100) to generate a distribution of the difference in proportions between the two groups.

P-values were calculated via the *boot*.*pval* function by inverting the bootstrap confidence intervals^60^. Under this approach, the p-value for the two-sided test is defined as the smallest significance level (alpha) such that the null hypothesis (a difference of zero) is excluded from the corresponding *1-alpha* percentile confidence interval.

### Recovery of a T-cell activation trajectory using diffusion maps

To recover T-cell activation trajectories from our Jurkat targeted sci-Plex single-cell profiles, we used diffusion maps, a spectral method for nonlinear dimensionality reduction^54,55^. Briefly, we used the diffusion map implementation in the *destiny* package in R^54,55^. The *destiny* implementation incorporates single-cell-specific features, including an efficient nearest-neighbor approximation, and optimizes the parameter selection for σ. The input for destiny consisted of the top 9 principal components as recommended for sparse scRNA-seq data^61^. Examination of the diffusion components (DCs), the eigenvectors of a Markovian transition probability matrix, identified DC1 as capturing the progression along T-cell activation when comparing (1) the distribution of controls to the T-cell activating exposure groups at different doses and (2) examining the relative levels of activation markers across DC1. We then used the ranked DC1 values to calculate a pseudotime measure.

### Grouping of T-cell activating exposures using multi-resolution variational inference (MrVI)

We used multi-resolution variational inference (MrVI)^56^ to group chemical exposures that modify T-cell activation by transcriptional similarity, as we have performed previously^18,20^. MrVI employs a hierarchical probabilistic framework that presumes a cell originates from nested experimental designs where individual cells are drawn from a sample, and samples are from experimental batches. The model learns two latent feature spaces: a sample-unaware *U-space*, which captures fundamental cell state features and is decoupled from the sample effects, and a sample-aware *Z-space*, which incorporates the effects of the sample while accounting for technical factors. MrVI (scvi-tools 1.4.0) uses multi-head cross-attention-based decoders to facilitate information flow and mapping features from U-space to Z-space.

We trained a MrVI model with the sample key defined as a unique exposure-dose combination. For all trained models, we used the recommended default model arguments in the scvi-tools implementation of MrVI with the treatment_dose combination corresponding to the sample_key and with max_epoch 150 which was sufficient given the validation evidence lower bound (ELBO). To select for gene features inputted into the MrVI models, we obtained the union of the probe genes and the 500 top highly variable genes (Scanpy) for the total dataset. This leverages the background genes. To visualize the output of the MrVI models, we generated hierarchically clustered heatmaps to group exposure-dose combinations.

### T-cell receptor detection with MixCR

T-cell receptor (TCR) sequences captured using TRTL in our targeted sci-Plex experiment were aligned, error-corrected, and assembled into clonotypes using the MiXCR software suite. The generic-lt-single-cell-amplicon-with-umi workflow was adapted in a custom .yaml file with input file name expansion using a custom whitelist of cell barcodes. Only reads with valid cell barcodes were retained for subsequent analyses, and TCR reads were assigned to individual cells. The custom pipeline steps include: raw paired-end sequencing reads were aligned to the human reference repertoire (V, D, J, C segments) using MiXCR *align*. Following alignment, UMI and cell tags were refined using *refineTagsandSort*. Clonotypes were then assembled with MiXCR *assemble*, which uses a quality-aware consensus sequence for the complementarity-determining region 3 (CDR3) for reads with the same UMI and cell barcode. Finally, the resultant clonotype tables were exported, and the VDJ gene hits were compared to the Jurkat E6 ground truth sequence^62^.

