## Supplementary Information for "A modular transcript enrichment strategy for scalable, atlas-aligned, and clonotype-resolved single-cell transcriptomics"

### Supplementary materials

Comparison of gene-targeted methods

| Application | Targeted sci-Plex | 10x Flex | Parse Gene Select | 10x Targeted Gene expression (discontinued) | HybriSeq |
|---|---|---|---|---|---|
| Targeted gene capture | low-cost probes adapted using TRTL for reverse transcription | low-cost probes used for PCR based counting | Biotinylated probes spanning whole transcript after whole transcriptome library preparation [1] | Biotinylated probes spanning whole transcript after whole transcriptome library preparation [2] | low-cost probes used for PCR based counting [3] |
| Whole transcript | yes | no | yes | yes | no |
| TCR/BCR profiling | yes | no | yes, using Evercode TCR | yes, if used with 5’ | no |
| CRISPR guide capture | yes | yes, with custom probes for each guide | yes using CRISPR Detect | yes | yes, in principle, with custom probes for each guide |
| Combinatorial indexing | yes | no | yes | no | yes |

Mode of TRTL integration

| | sci-Plex | Split-seq/ Parse | PERFF-seq |
|---|---|---|---|
| TRTL integration | yes: tested | yes: replace first round RT using dT-barcode1 primers with TRTL | hybridization chain reaction (HCR) with TRTL-ON buffer conditions |

**Supplementary Figure 1: Comparison of gene-targeted methods and compatibility of TRTL, in principle, with single-cell methodologies** (a) comparison of applications of existing gene-targeting methods (b) strategies for TRTL integration with other single-cell RNA profiling methods. Abbreviations: polymerase chain reaction (PCR), CRISPR (Clustered Regularly Interspaced Short Palindromic Repeats), capture sequence 1 (CS1)[10], unique molecular identifier (UMI).

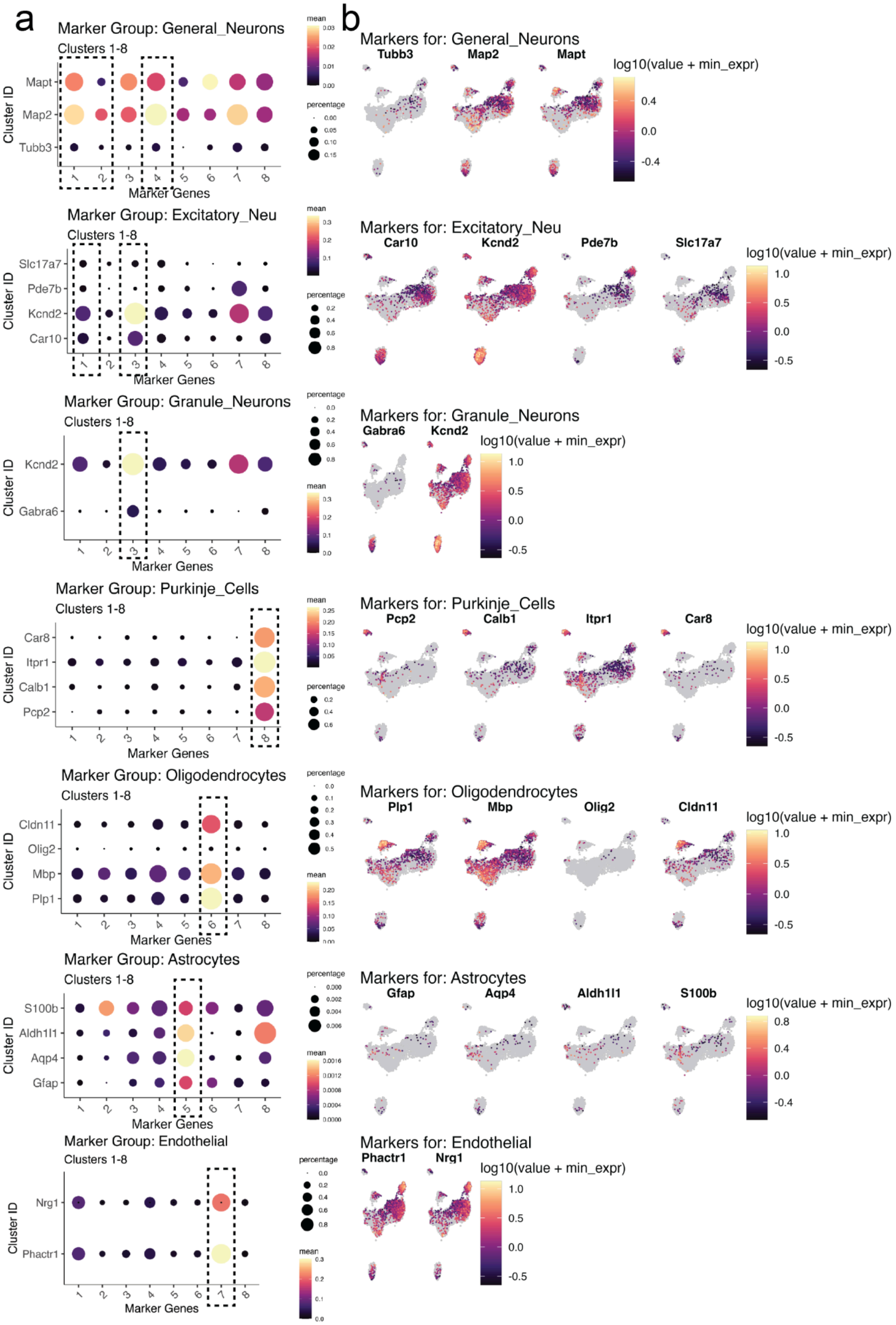
a
b
Marker Group: General_Neurons
Marker Group: Excitatory_Neu
Marker Group: Granule_Neurons
Marker Group: Purkinje_Cells
Marker Group: Oligodendrocytes
Marker Group: Astrocytes
Marker Group: Endothelial
Clusters 1-8
Cluster ID
Marker Genes
mean
percentage
Markers for: General_Neurons
Tubb3
Map2
Mapt
Markers for: Excitatory_Neu
Car10
Kcnd2
Pde7b
Slc17a7
Markers for: Granule_Neurons
Gabra6
Markers for: Purkinje_Cells
Pcp2
Calb1
Itpr1
Car8
Markers for: Oligodendrocytes
Plp1
Mbp
Olig2
Cldn11
Markers for: Astrocytes
Gfap
Aqp4
Aldh1l1
S100b
Markers for: Endothelial
Phactr1
Nrg1
log10(value + min_expr)

**Supplementary Figure 2: Cell type and cell lineage specific marker expression in mouse brain samples subjected to transcriptome-wide combinatorial indexing single- cell RNA-seq profiling.** (a) Dot plot summarizing the expression of canonical marker genes across Leiden-based clusters from **Figure 1c**. The color intensity represents the average expression level within each cluster, while the dot size indicates the percentage of cells expressing the gene. (b) UMAP embeddings as in **Figure 1c** colored by the log-normalized expression of select marker genes from (a).

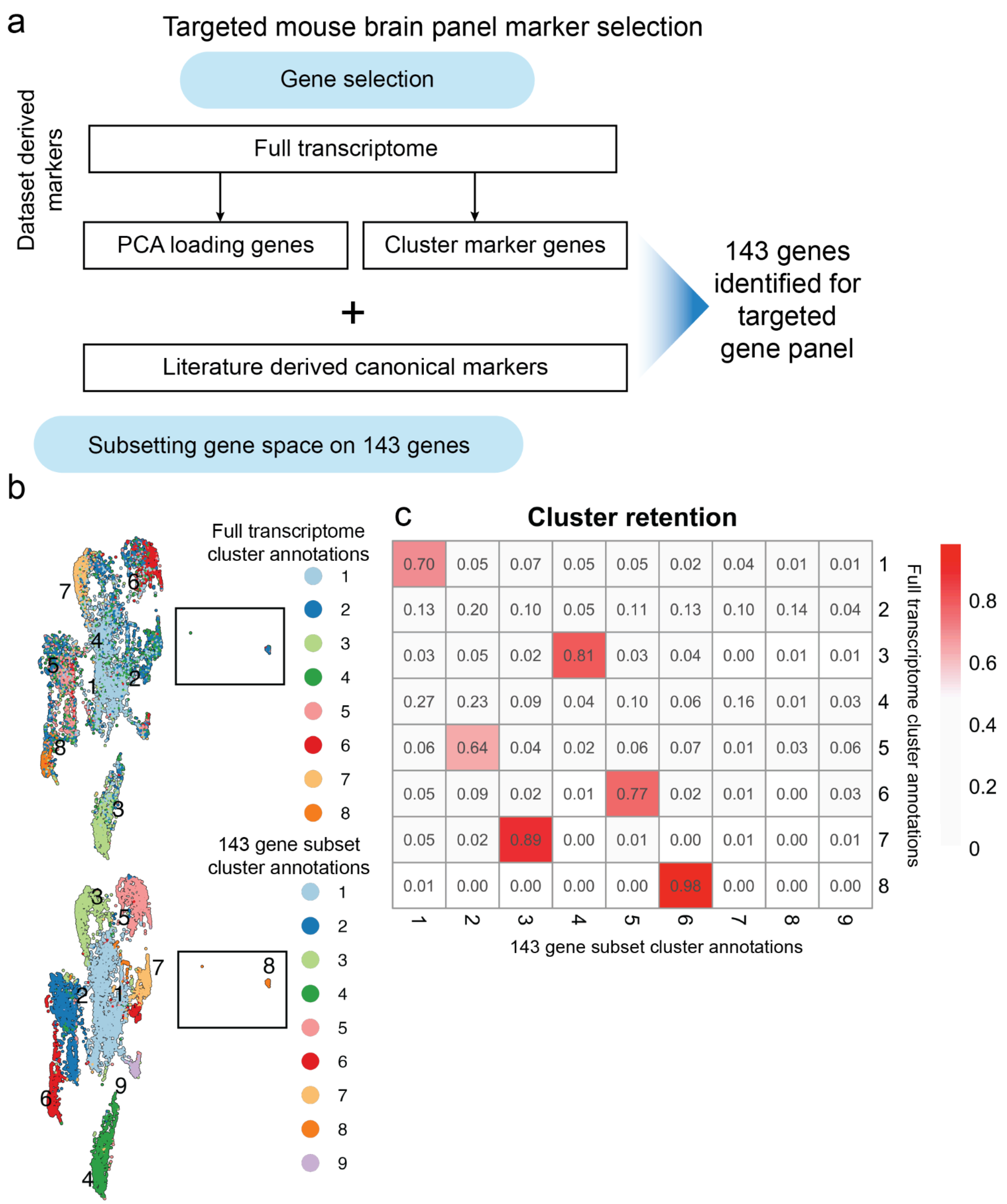


**Supplementary Figure 3: Mouse brain panel selection** (a) Schematic of our approach to prioritize a 143-target gene panel for targeted profiling of the mouse brain. (b) UMAP embedding of cells from [main figure call out] created using the subset of 143-target genes as feature for dimensionality reduction. Top: color refers to the cluster assignment for each cells as identified using the full transcriptome as in **Figure 1d**. Bottom: Cells colored by cluster identified from Leiden-based community detection after dimensionality reduction on the 143-target gene subset. (c) Heatmap depicting the proportion of cells from clusters identified using the full transcriptome (rows) or across the dimensionality reduction obtained across the 143-target gene subset (columns)

**a** Mouse brain dT vs TRTL experimental overview

| | Mouse brain experiments | # of fresh frozen brains processed |
|---|---|---|
| Experiment 1 | Comparison of **dT sci-RNA-seq3 to TRTL** | 1 |
| Experiment 2 | Comparison of **dT sci-RNA-seq3, TRTL, and TRTL-ON** | 2 |

**b** cells above cutoff:

22226 cells above cutoff dT

46809 cells above cutoff TRTL

21715 cells above cutoff TRTL-ON

**c**

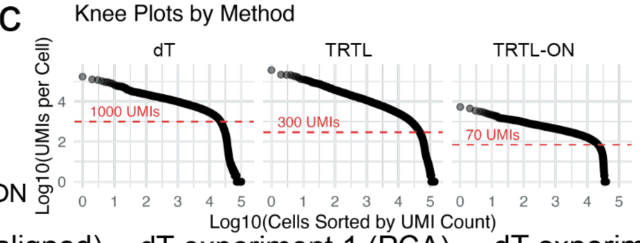


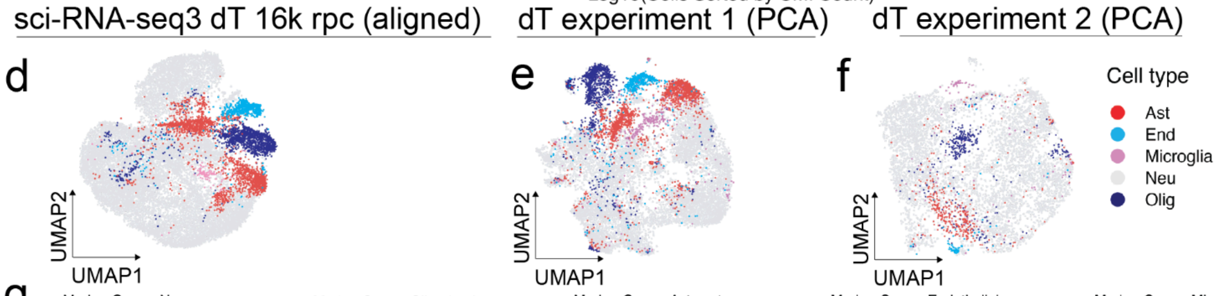


**g**

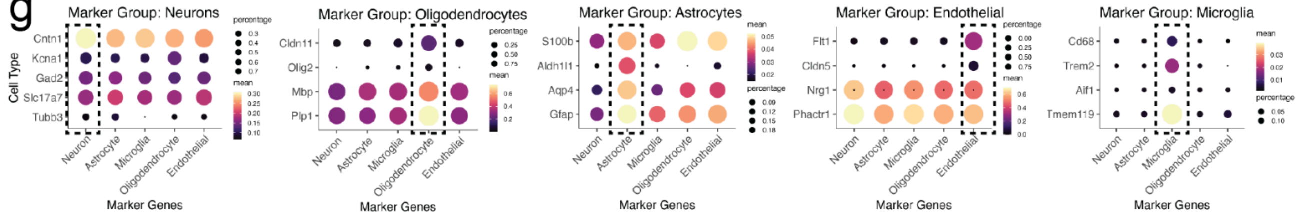


**h** Dropout of probe genes in TRTL-ON

| Experiment | Probe Genes Detected (> 1 Count) | Probe Genes Detected (> 1 Count) in at least 0.5% of total cells | Median genes expressed |
|---|---|---|---|
| **TRTL-ON** | 141 | 85 | 142 |
| **dT 16k** | 143 | 127 | 1939 |
| **dT 7k** | 142 | 121 | 1603 |
| **TRTL 7k** | 143 | 120 | 754 |

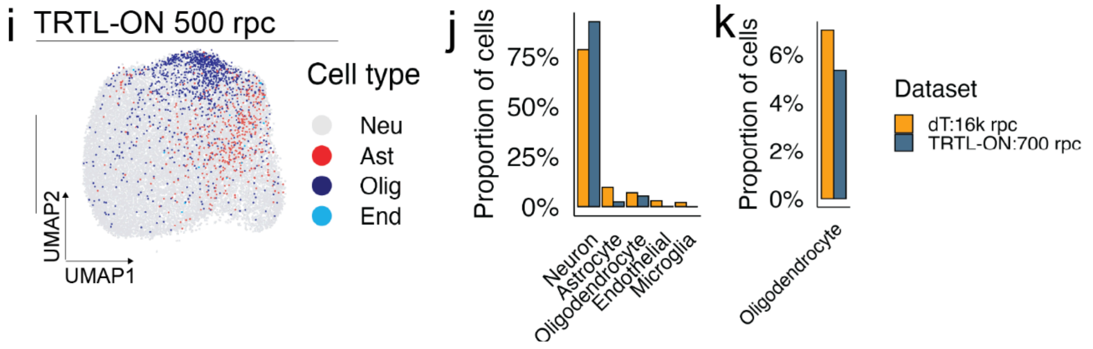

**Supplementary Figure 4: Transcriptome-wide oligo-dT vs TRTL and TRTL-ON sci-RNA-seq profiling(a)** Overview of transcriptome-wide sci-RNA-seq, TRTL and TRTL-ON experiments **(b)** Cell numbers after filtering each experimental condition (dT, TRTL, and TRTL-ON) by cellular transcriptome quality as measured by distribution of UMI counts. **(c)** Knee-plots for each experimental condition with cutoffs used to define cells in **(b). (d)** Batch aligned UMAP embedding of transcriptome-wide oligo-dT sci-RNA-seq. Batches were aligned using the mutual nearest neighbor (MNN) approach on the top principal component of each batch. Color denotes cell type assignment. **(e-f)** UMAP embedding of transcriptome-wide oligo-dT sci-RNA-seq3 experiments 1 (left) and 2 (right) without batch alignment. Color denotes cell type assignments **(g)** Dot plot summarizing the expression of canonical marker genes across Leiden-based cluster cell-type assignments from **b**. The color intensity represents the average expression level within each cluster, while the dot size indicates the percentage of cells expressing the gene. **(h)** Probe gene detection across full and downsampled TRTL and dT datasets. **(i)** UMAP embedding of single-cell transcriptomes obtained using our TRTL-ON approach. Color denotes cell type classification using  Seurat's PCA-based anchor label transfer from dT 16k rpc to TRTL-ON **(j-k)** Cell-type proportions annotated for dT and TRTL-ON datasets after mapping to the full transcriptome-wide reference generated by our sci-RNA-seq3 protocol. (k) Oligodendrocyte proportions as in (j) in the TRTL-ON condition.

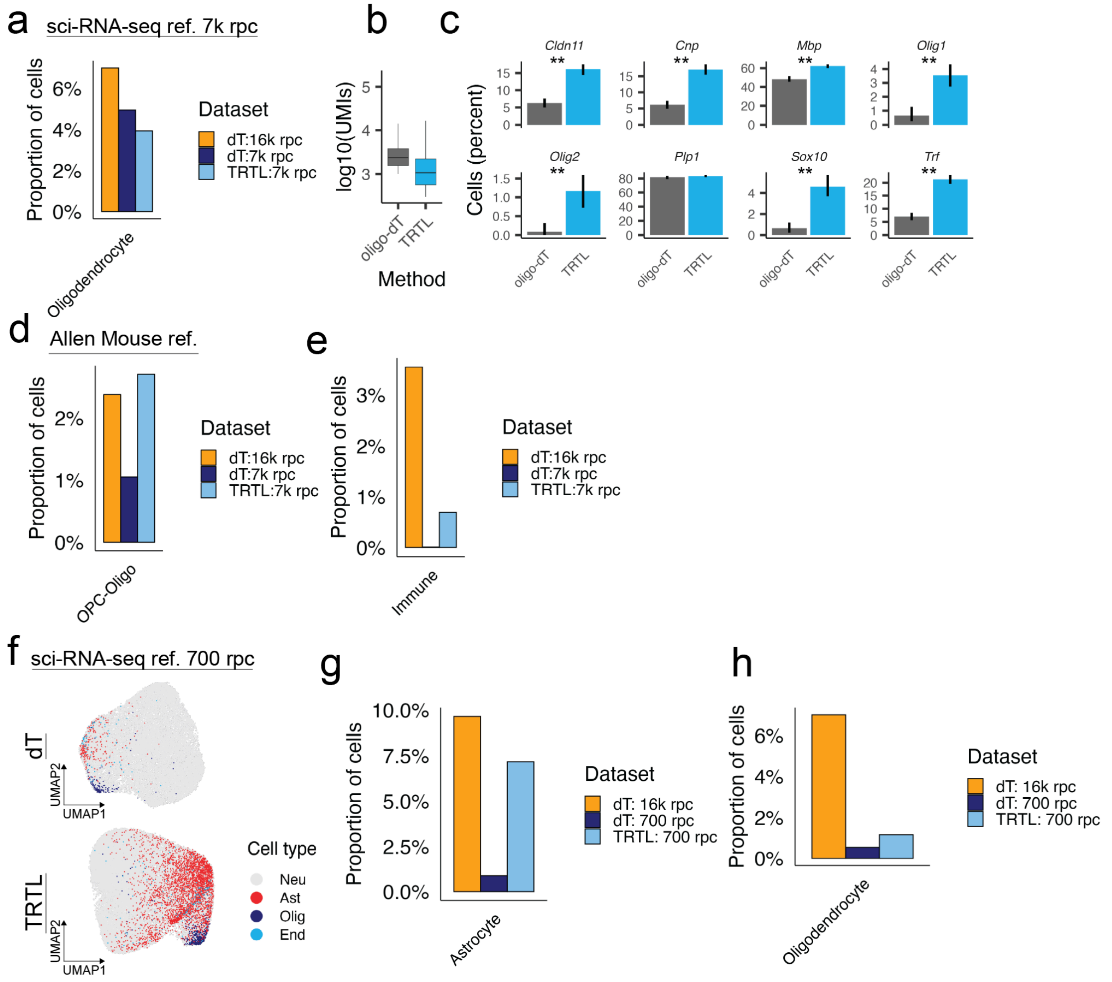


**Supplementary Figure 5: TRTL 7k and 700 reads per cell (rpc).** (**a**) Oligodendrocyte proportions annotated for downsampled dT and TRTL datasets at 7k rpc mapped to the full transcriptome-wide reference generated by our sci-RNA-seq3 protocol. (**b**) UMI counts for oligodendrocyte assigned cells in the downsampled 7k rpc datasets (**c**) Percent of cells expressing more than one transcript for oligodendrocyte marker genes in a dataset with equal information content per cell (normalized for UMI counts). Error bars indicate the 95% bootstrap confidence interval. Statistical significance of the difference between groups was determined by a non-parametric bootstrap test * $p < 0.05$, ** $p \leq 0.01$.(**d**) OPC-Oligo proportions as in (**a**) mapped to the Allen Mouse Brain Atlas. (**e**) Immune cell proportions as in (**a**) mapped to the Allen Mouse Brain Atlas. (**f**) Batch aligned UMAP embedding of downsampled (700 reads per cell [rpc]) for transcriptome-wide dT (top) and TRTL targeted (bottom) datasets. Color denotes the cell type classification using Seurat's PCA-based anchor label transfer from the full transcriptome-wide reference generated by our sci-RNA-seq3 protocol. (**g-h**) Astrocyte (g) and oligodendrocyte (h) proportions annotated for downsampled dT and TRTL datasets at 700 rpc mapped to the full transcriptome-wide reference generated by our sci-RNA-seq3 protocol.

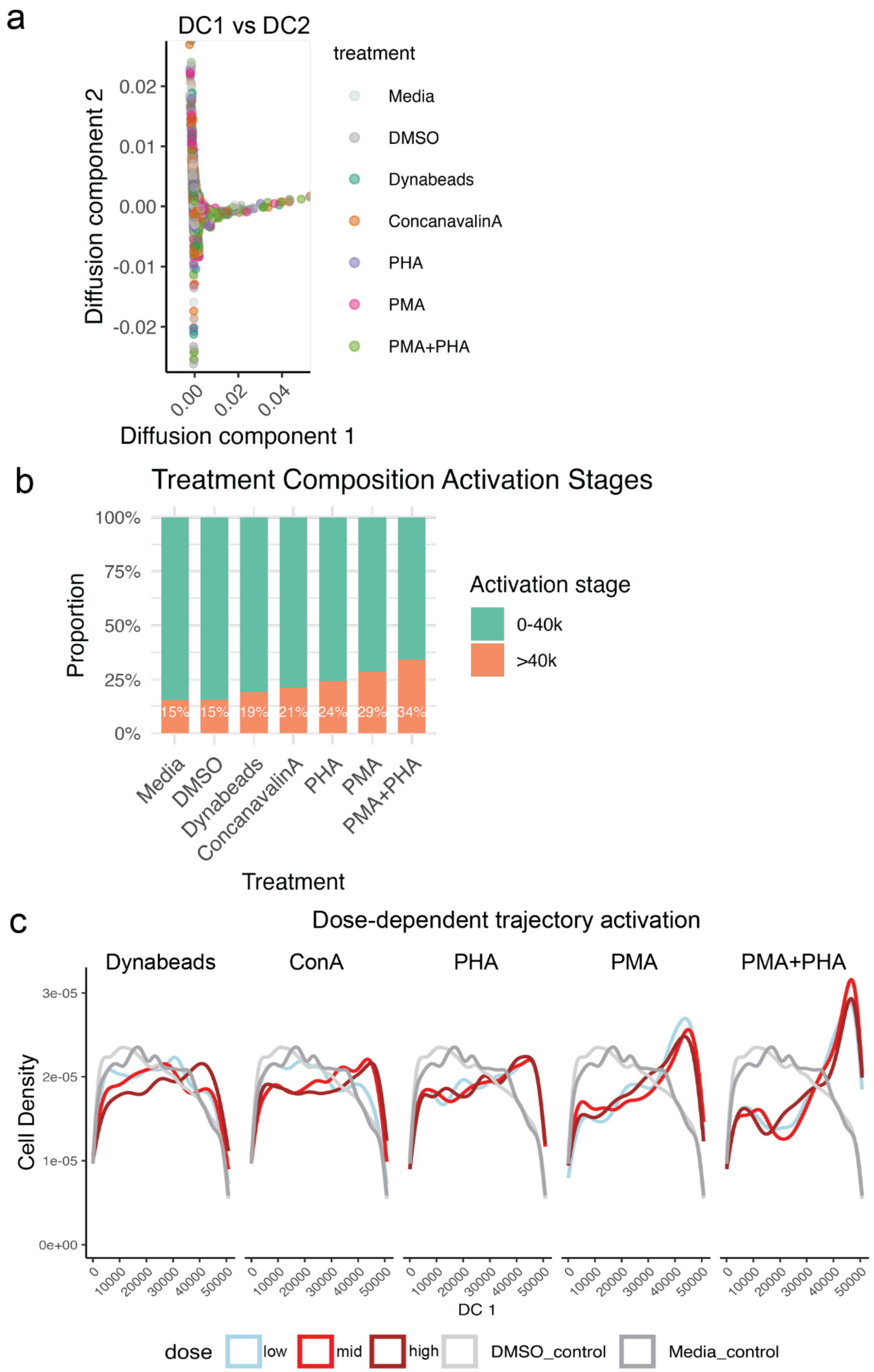
a
DC1 vs DC2
Diffusion component 2
0.02
0.01
0.00
-0.01
-0.02
0.00
0.02
0.04
Diffusion component 1
treatment
Media
DMSO
Dynabeads
ConcanavalinA
PHA
PMA
PMA+PHA
b
Treatment Composition Activation Stages
Proportion
100%
75%
50%
25%
0%
15%
15%
19%
21%
24%
29%
34%
Media
DMSO
Dynabeads
ConcanavalinA
PHA
PMA
PMA+PHA
Treatment
Activation stage
0-40k
>40k
c
Dose-dependent trajectory activation
Dynabeads
ConA
PHA
PMA
PMA+PHA
Cell Density
3e-05
2e-05
1e-05
0e+00
0
10000
20000
30000
40000
50000
DC 1
dose
low
mid
high
DMSO_control
Media_control

**Supplementary Figure 6: Recreating a continuous T-cell activation trajectory with diffusion maps. (a)** Dimensionality reduction of targeted-sci-Plex T-cell activation single-cell data using diffusion maps across the top two diffusion components (DC1 and DC2). Each cell is colored by the condition to which it was exposed. **(b)** Proportion of cells that reach the end of the T-cell activation trajectory defined as progression along DC1 (pseudotime value > 40,000) by treatment. **(c)** Density of cells across the activation trajectory defined as progression along DC1 by treatment and dose (low/mid/high). Low, medium, and high doses respectively: Dynabeads (1, 2, 4 µL), ConA (1.25, 2.5, 5 µg/mL), PHA (1.25, 2.5, 5 µg/mL), PMA (50, 100, 200 ng/mL), and PHA+PMA (1.25 µg/mL + 50 ng/mL, 2.5 µg/mL + 100 ng/mL, 5 µg/mL + 200 ng/mL). DMSO and media only controls are in different shades of grey.

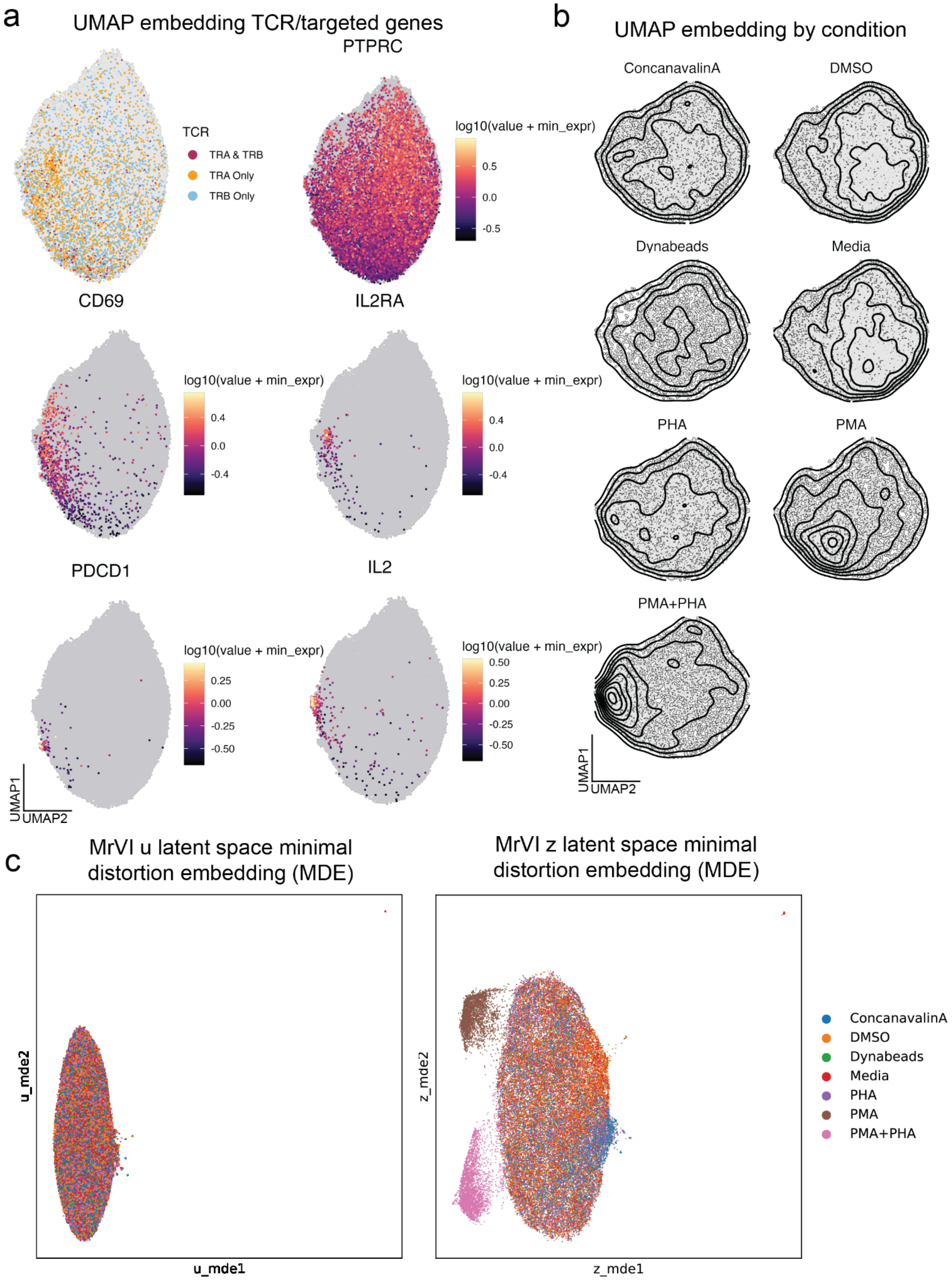

a
UMAP embedding TCR/targeted genes
PTPRC
TCR
TRA & TRB
TRA Only
TRB Only
log10(value + min_expr)
0.5
0.0
-0.5
CD69
IL2RA
log10(value + min_expr)
0.4
0.0
-0.4
log10(value + min_expr)
0.4
0.0
-0.4
PDCD1
IL2
log10(value + min_expr)
0.25
0.00
-0.25
-0.50
log10(value + min_expr)
0.50
0.25
0.00
-0.25
-0.50
UMAP1
UMAP2
b
UMAP embedding by condition
ConcanavalinA
DMSO
Dynabeads
Media
PHA
PMA
PMA+PHA
UMAP1
UMAP2
c
MrVI u latent space minimal distortion embedding (MDE)
MrVI z latent space minimal distortion embedding (MDE)
u_mde2
u_mde1
z_mde2
z_mde1
ConcanavalinA
DMSO
Dynabeads
Media
PHA
PMA
PMA+PHA

**Supplementary Figure 7: Deep learning based classification of T-cell activating exposures from a targeted-sci-Plex readout.** **(a)** UMAP embeddings of single-cell transcriptomes obtained from a targeted-sci-Plex screen of T-cell activation conditions colored by the specified gene captured per cell. **(b)** UMAP embeddings as in A with the density of cells treated with each exposure overlayed. **(c)** MrVI minimal distortion embeddings for the sample unaware u (left) and sample aware Z space. Cells are colored by the treatment to which a cell was exposed.

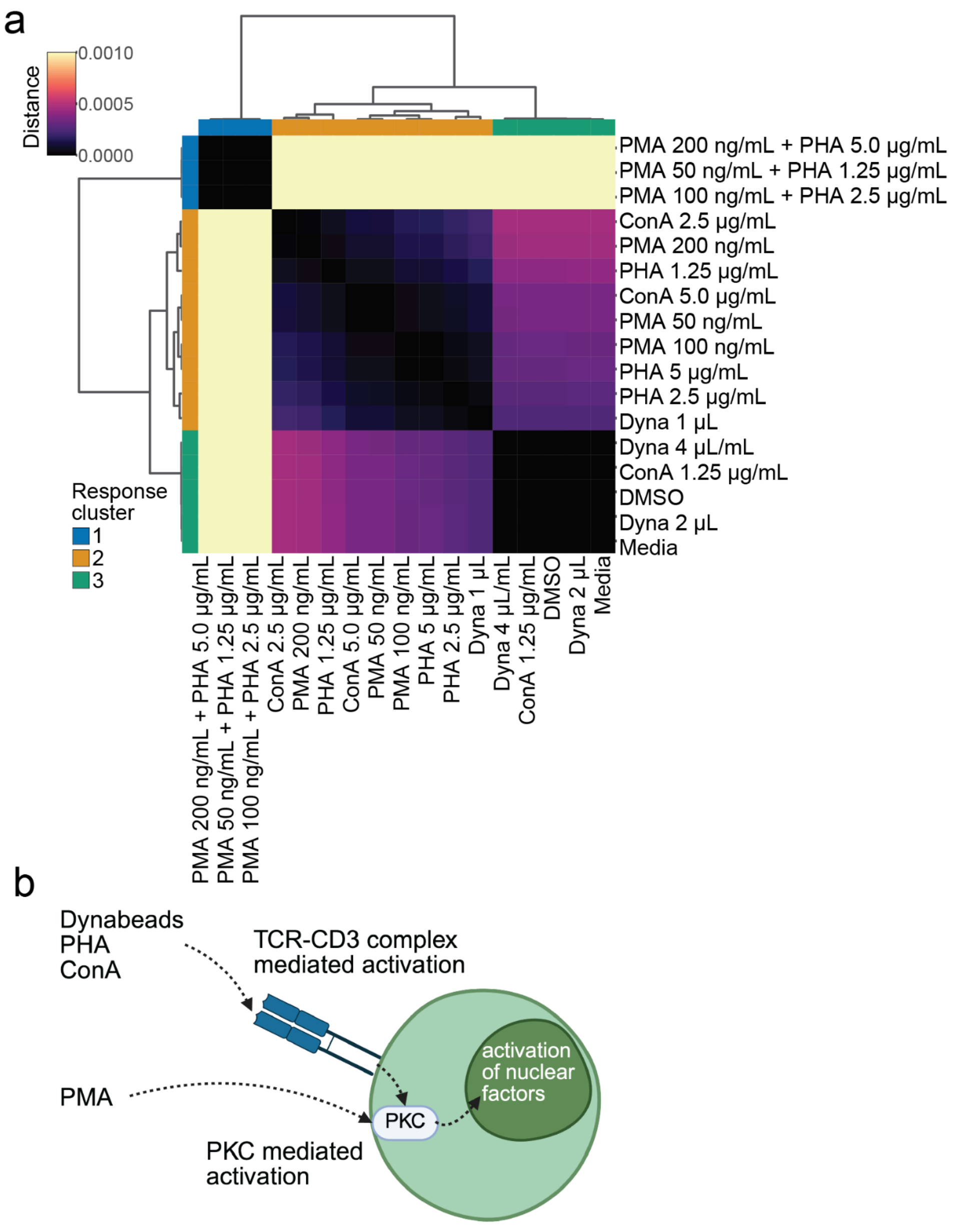


**Supplementary Figure 8: Classification of activating agents based on only probe genes** **(a)** Heatmap of the mean transcriptional distances between treatment conditions, estimated with MrVI using only probe genes as the input. Hierarchical clustering of the pairwise distances reveal three distinct treatment clades based on transcriptional similarity and activation state. **(b)** Diagram of activation pathways for Dynabeads, PHA, ConA (TCR-CD3 mediated activation via PKC) compared to PMA (PKC mediated activation).

**Supplementary Table 1: Probes and oligonucleotides used in this study (provided as a supplemenatry xlsx file)** final mouse brain panel, number of probes per gene in the mouse brain studies, final Jurkat panel, number of probes per gene in the Jurkat studies, oligos used in studies

| Experiment: condition | Total cells (after filtering) | Median UMIs per cell | Average reads per cell (estimated) | Duplication rate |
|---|---|---|---|---|
| 1: dt | 15,395 | 3,377 | 16k | 31.3% |
| 1: TRTL | 21,967 | 1,744 | 16k | 35.4% |
| 2: dt | 6,831 | 3,372 | 15k | 21.0% |
| 2: TRTL | 24,842 | 1,128 | 11k | 30.7% |
| 2: TRTL-ON | 21,715 | 155 | 1k | 29.7% |

Results of random sampling to 7000 rpc

| Experiment: condition | Reads (aligned) | Average reads per cell (aligned) | Duplication rate |
|---|---|---|---|
| 1: dt | 148,056,218 | 9,617 | 18.2% |
| 1: TRTL | 167,756,845 | 6,752 | 26.4% |
| 2: dt | 50,390,346 | 7,376 | 14.9% |
| 2: TRTL | 191,803,666 | 7,720 | 30.7% |

Results of random sampling to 700 rpc

| | | | |
|---|---|---|---|
| 1: dt | 12,189,350 | 792 | 2.7% |
| 1: TRTL | 16,090,616 | 648 | 3.4% |
| 2: dt | 4,652,116 | 681 | 1.6% |
| 2: TRTL | 15,343,216 | 698 | 3.4% |

**Supplementary Table 2: Overall sequencing and subsampling summary for mouse brain**
